# Unsupervised learning elucidates the interplay between conformational flexibility and aggregation in synergistic antimicrobial peptides

**DOI:** 10.1101/2024.07.03.601908

**Authors:** Miruna Serian, A. James Mason, Christian D. Lorenz

## Abstract

Synergy between antimicrobial peptides (AMPs) may be the key to their evolutionary success and could be exploited to develop more potent antibacterial agents. One of the factors thought to be essential for AMP potency is their conformational flexibility, but characterizing the diverse conformational states of AMPs experimentally remains challenging. Here we introduce a method for characterising the conformational flexibility of AMPs and provide new insights into how the interplay between conformation and aggregation in synergistic AMP combinations yields emergent properties. We use unsupervised learning and molecular dynamics simulations to show that mixing two AMPs from the Winter Flounder family (pleurocidin (WF2) & WF1a) constrains their conformational space, reducing the number of distinct conformations adopted by the peptides, most notably for WF2. The aggregation behaviour of the peptides is also altered, favouring the formation of higher-order aggregates upon mixing. Critically, the interaction between WF1a and WF2 influences the distribution of WF2 conformations within aggregates, revealing how WF1a can modulate WF2 behaviour. Our work paves the way for deeper understanding of the synergy between AMPs, a fundamental process in nature.

## Introduction

Antimicrobial resistance (AMR) is one of the most significant global public health threats. Despite this, the development of new antibiotics has declined, and the World Health Organization (WHO) describes the antibacterial clinical and preclinical pipeline as stagnant and far from meeting global needs (1). Therefore, there is an urgent need for increased efforts to develop alternatives to current antimicrobial agents. Consequently, the generation of novel medications to control and treat infections caused by multidrug-resistant pathogens has become a pressing priority for the scientific community. Antimicrobial peptides have quickly gained traction as promising drug candidates because of their potency against both Gram-negative and Gram-positive bacteria (2). Antimicrobial peptides are evolutionary conserved components of the immune system, found in almost all life forms, from prokaryotes to humans (3), and have been shown to have distinct roles. While in higher life forms they are produced to protect the host against infection, bacteria can also produce AMPs to kill other bacteria competing for the same environment (4). Despite their potential, several challenges hinder the widespread use of antimicrobial peptides as antibiotics alternatives. These include concerns about host toxicity (5), the emergence of bacterial resistance (6) and high production costs (7). To address these limitations, combining different antimicrobial peptides has emerged as a promising strategy. Similar to combination therapy with traditional drugs, combining antimicrobial peptides can lead to synergistic effects, potentially reducing the necessary dosage, minimizing side effects, and lowering the risk of resistance development (8).

Despite the benefits of synergistic combinations of AMPs, the mechanisms of their synergy are not yet fully understood. Computational and experimental studies have proposed several mechanisms, including pore formation. For instance, combining PGLa and MAG2, two peptides produced in the skin of *Xenopus laevis* can lead to the formation of a toroidal pore structure, that can in turn lead to more membrane disruption (9). In other cases, such as the interaction between the two AMPs produced by bumblebees, abaecin and the pore forming AMP hymenoptaecin, distinct mechanisms are observed; hymenoptaecin forms membrane pores, destabilizing bacterial membranes and allowing abaecin entry into bacterial cells (10). Abaecin can also synergise with pore-forming peptides from other organisms (11). Additionally, synergistic behaviour may arise from complementary mechanisms, such as the in the case of coleoptericin and defensin. Coleoptericin acts to improve host survival while defensin can reduce bacterial load (12). Overall, pore formation or peptide aggregation remains among the most suggested mechanisms of action for synergistic peptides.

Further, there is a scarcity of research concerning the synergy between antimicrobial peptides originating from the same species. Previous work has investigated the synergy between the family of Winter Flounder (WF) peptides employing an interdisciplinary approach of microbiology, biophysics and electrophysiology (13). Despite only two out of the six WF peptides exhibiting potent antimicrobial activity when used individually, the study identified a series of two-way combinations that were active against both Gram-positive and Gram-negative bacteria.

The Winter Flounder peptides are a family of peptides extracted from the *Pseudopleuronectes americanus* fish and they are found in the gills and intestines, as well as the skin of the fish (14–16). Among the six peptides, pleurocidin (or WF2) is the most studied peptide due to its broad spectrum antimicrobial activity (17–20). Pleurocidin functions through a combination of membrane destabilization and metabolic inhibition. While all six WF peptides share some sequence similarity, the majority exhibit limited efficacy against bacteria, except for pleurocidin.

Further, the conformational flexibility of AMPs is considered to be crucial for their antibacterial potency (20–22). For example, in the case of magainin, reduced flexibility caused by the cyclization of the peptide, results in reduced antimicrobial activity (23). Other studies showed that the conformational flexibility of *α*-helical antimicrobial peptides determines their anti-cancer and antimicrobial potency (21, 22). Therefore, we expect that conformation plays an important role in the synergistic activity of AMPs. However, it remains challenging to isolate and characterize the different conformational states adopted by AMPs experimentally, which is further exacerbated by the heterogeneity observed in AMPs systems (13). Therefore, we employed a combination of unsupervised machine learning and molecular dynamics simulations to investigate the interplay between AMPs aggregation and conformation in a pair of synergistic AMPs from the Winter Flounder family, in a model of Gram-positive bacteria. Previous research revealed that WF peptides could assemble into dimers, which was observed during 200ns molecular dynamics simulations (13). Our new findings reveal an interplay between the conformational flexibility of the WF peptides and the ability to form higher order aggregates when the peptides are simulated for 1 microsecond.

## Methods

### Peptide-Lipid Systems

The starting structures of the WF peptides were obtained from PDB (24), and were resolved using NMR in SDS-d_25_ micelles. The PDB entries contained 100 conformers, out of which the conformer with the lowest potential energy was selected as the representative structure for each respective peptide. The systems were constructed using eight peptides placed 20 Å above a lipid bilayer in a random position and orientation. The pure systems contain eight peptides of the respective type, while the combination systems contain four peptides of each type, with a total of eight peptides. The membrane used in the simulations is made of 256 POPG lipids and was generated using CHARMM-GUI Membrane Builder (25) using the CHARMM36 parameters. The peptide to lipid ratio is 1:32.

#### A. Molecular Dynamics Simulations

Simulations were run using the GROMACS package (26). Periodic boundary conditions were applied in all directions. TIP3P model was used to describe the water that solvated the system. The final system contains Na+ ions in order to insure that the overall charge of the system is 0. The system were energy-minimised at 310 K with the Nose-Hoover Thermostat using the steepest descent algorithm to remove steric clashes between atoms or any artificially large energy. The density of the system was then equilibrated under the isothermal-isobaric NPT ensemble for 2 ns using a Berendsen Thermostat at 310 K. Production simulations were run for 1 *µ*s. The leapfrog algorithm and a 2 fs time step were used to integrate the equations of motion in all of the simulations. A cutoff of 1.2 nm was applied for van der Waals and electrostatic interactions using the Particle-Mesh Ewald algorithm for the long range electrostatic interactions. Hydrogen containing bonds were constrained using the LINCS algorithm. The systems were simulated in two replicas.

#### B. Aggregation and Intermolecular Interactions Analysis

The resulting data from the molecular dynamics simulations was analysed using in-house scripts written in python and taking advantage of the MDanalysis package (27). Only the last 500 ns of the simulations were used for all the analyses presented in this work. Data from both the initial and replicate runs for each system were included in the calculations.

The peptides were considered to be in an aggregate if the minimum distance between any two residues on each peptide was *d ≤* 6 Å. The aggregates were identified using graph theory and the NetworkX library (28).

#### C. UMAP - HDBSCAN Clustering

Clustering was performed using UMAP (29) and HDBSCAN (30) on the last 500 ns of the simulations using the n-n distances between all C*α* atoms of each residue. The n-n distance data from both replicates, for both the pure and mixed systems, served as input for the clustering workflow. The hyperparameters for UMAP and HDBSCAN were selected iteratively for each type of peptide and can be found in Table 1. The structural meaningfulness of the resulting clusters was tested by analysing their structural properties, including the distribution of the RMSD and radius of gyration values for each conformational cluster. This methodology was applied individually to each type of peptide. More details about how we implemented the methods can be found in Fig. SI 1. Previously, we have used a similar approach to analyse the conformations taken by polymers (31–34).

**Table 1.**
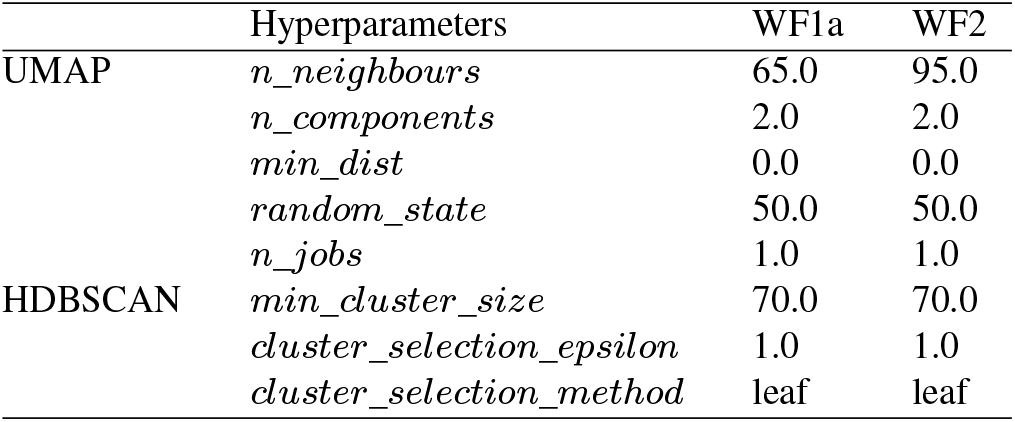
Hyperparameters used for UMAP and HDBSCAN.

|  | Hyperparameters | WF1a | WF2 |
| --- | --- | --- | --- |
| UMAP | *n_neighbours* | 65.0 | 95.0 |
|  | *n_components* | 2.0 | 2.0 |
|  | *min_dist* | 0.0 | 0.0 |
|  | *random_state* | 50.0 | 50.0 |
|  | *n_jobs* | 1.0 | 1.0 |
| HDBSCAN | *min_cluster_size* | 70.0 | 70.0 |
|  | *cluster_selection_epsilon* | 1.0 | 1.0 |
|  | *cluster_selection_method* | leaf | leaf |

The representative structure for each conformational cluster was selected as the structure with the RMSD value closest to the mean RMSD of all conformations within that cluster. The reference structure used for calculating the RMSD values was the same corresponding peptide structure (WF1a or WF2) in the mixed system. Secondary structures for each cluster were computed using the DSSP package, based on the work of Kabsch and Sander (35). The eight different secondary assignments computed by DSSP are described in Table 2. Conformation snapshots were generated using VMD (36).

**Table 2.** DSSP Dictionary.

| Letter | Secondary Structure |
| --- | --- |
| H | α-helix |
| B | Residue in isolated β-bridge |
| E | Extended strand, participates in β-ladder |
| G | $3_{10}$-helix |
| I | π-helix |
| T | Hydrogen bonded turn |
| S | Bend |

#### D. Effects on the membrane lipids

The degree of insertion into the membrane was computed using the MDAnalysis package (27) and in-house python scripts. The insertion of the peptides into the membrane is determined by finding the mean *z*-positions of the amino acid residues within each peptide, relative to the plane of the phosphorous atoms in the lipid headgroups which compose the nearest leaflet of the lipid bilayer. The insertion per peptide is determined by the residue with the deepest insertion into the membrane. The area per lipid (APL) of the membrane was computed using in-house scripts via a 2D Voronoi tessellation (37). Only the APL values during the last 50 ns of the simulation for each system were analysed.

The membrane curvature was measured for each system and replica run for the upper leaflet only. The values were computed using the LipidDyn package (38). Negative values correspond to negative curvatures while positive values indicate positive curvatures.

## Results

### E. WF1a and WF2 combination leads to the formation of higher-order aggregates

We investigated the interaction between the peptides in two different systems, in which the peptides were placed on top of a model representative of Gram-positive bacteria composed of 256 POPG lipids. The two systems are split into pure systems, in which 4 peptides of each type were simulated individually, and the mixed systems, where the two peptides were simulated together in equal concentrations (4 peptides of each type, for a total of 8 peptides).

Our results suggest that both AMPs exhibit a propensity to interact with the other peptides in the system (Fig. 1 a), which has previously been reported (13). In the mixed systems, both WF1a and WF2 peptides exhibit a gradual increase in their aggregation probability, after which WF1a peptides show a higher probability of being found in aggregates than WF2 peptides. This indicates that most of the four WF1a peptides involved in the mixed systems are likely to be found in aggregates. When simulated individually, the two AMPs exhibit similar propensities to aggregate, with more than 50% of the peptides found in aggregates. Interestingly, mixing the two peptides increases the probability of WF1a peptides being involved in aggregates, while that of WF2 peptides is not greatly affected.

**Fig. 1.**
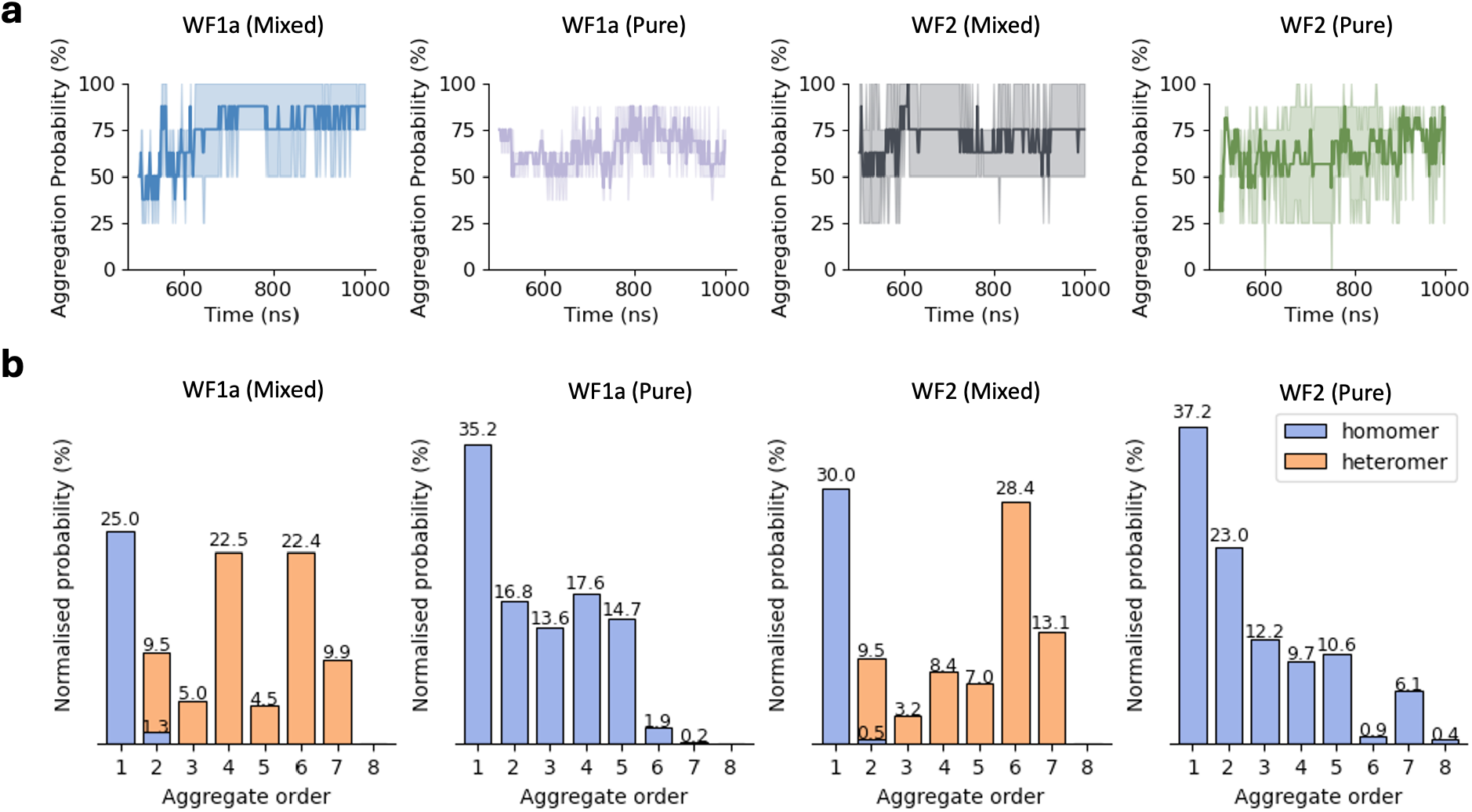
Peptide aggregation probabilities. (a) The aggregation probability was calculated across all both types of systems and include both simulation runs. The values were normalized by the number of peptides of the same type in the system. The resulting values indicate the likelihood of a specific peptide type being present within an aggregate at any given time. (d) Distribution of aggregate order probabilities is depicted for each peptide type and system, illustrating the likelihood of being in an aggregate of a specific size.

The impact of combining the two peptides on their aggregation preferences becomes apparent when examining the distribution of different orders of aggregates (Fig. 1 b). When simulated individually, both peptides predominantly exist as monomers, with similar occurrences of dimers and trimers. We observed more occurrences of 4-mers of WF1a peptides than WF2 peptides, which also demonstrated a limited capacity to form short-lived 8-mers. When considering the size of aggregates that are formed, we find that in the pure systems both peptides primarily form relatively small (*≤*5 peptides) aggregates. However, combining the two types of peptides induces a shift towards the formation of higher-order aggregates, such as 6-mers and 7-mers, which would not be expected purely based on chance. Despite the emergence of these larger aggregates, a substantial proportion of both peptide types still maintains a preference for existing as monomers, remaining unbound to other peptides.

In the mixed systems WF2 peptides are more prone to forming higher-order aggregates, such as 6-mers and 7-mers when compared to WF1a peptides (Fig. 1 b). This preference is also shown in our analysis of the composition of higher-order aggregates (*n ≥*3), which indicates that 3-mers and 4-mers comprise a greater proportion of WF1a peptides, whereas 5-mers, 6-mers, and 7-mers are more likely to contain a higher ratio of WF2 peptide (Table 3).

**Table 3.**
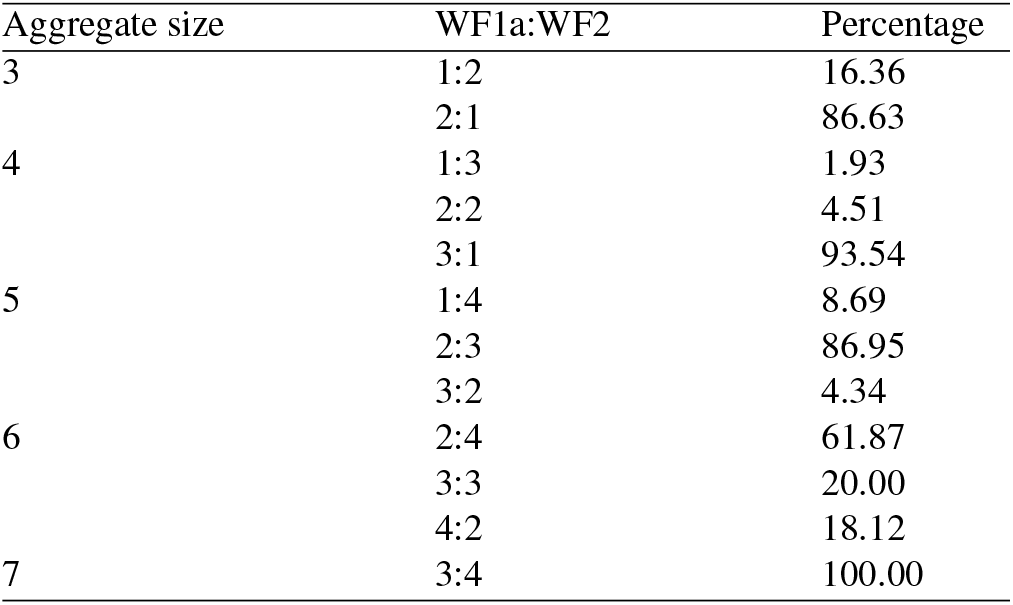
WF1a:WF2 peptide ratio occurrences in higher-order aggregat.

| Aggregate size | WF1a:WF2 | Percentage |
| --- | --- | --- |
| 3 | 1:2 | 16.36 |
|  | 2:1 | 86.63 |
| 4 | 1:3 | 1.93 |
|  | 2:2 | 4.51 |
|  | 3:1 | 93.54 |
| 5 | 1:4 | 8.69 |
|  | 2:3 | 86.95 |
|  | 3:2 | 4.34 |
| 6 | 2:4 | 61.87 |
|  | 3:3 | 20.00 |
|  | 4:2 | 18.12 |
| 7 | 3:4 | 100.00 |

**Table 4.**
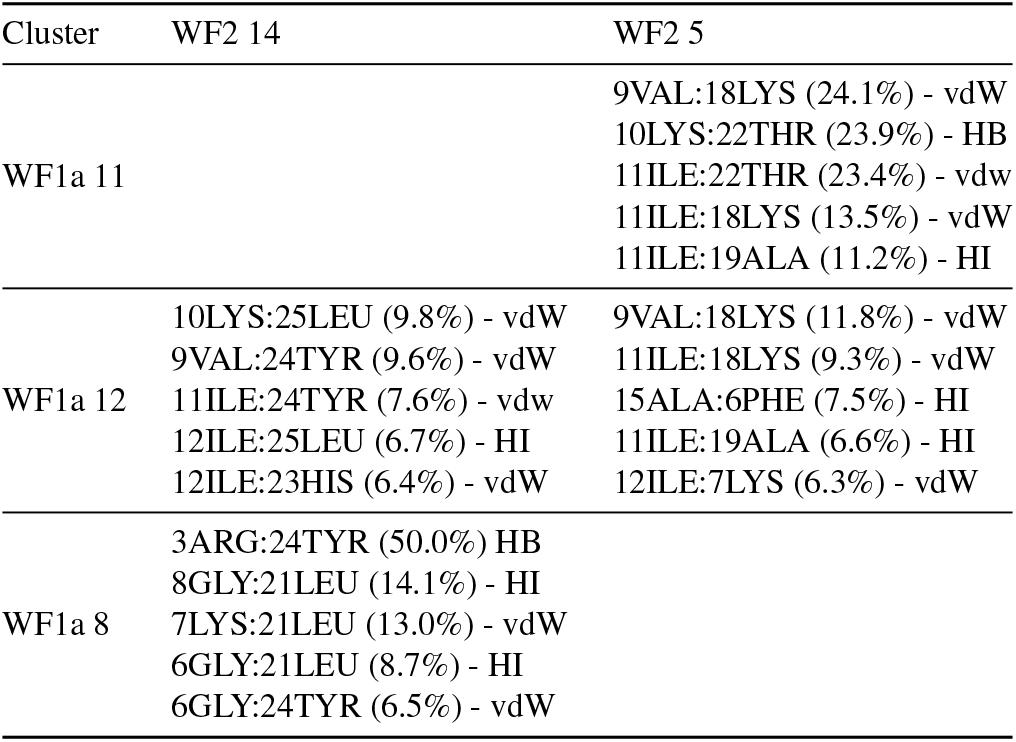
Top residue-residue contacts for WF1a-WF2 interactions.

### WF1a and WF2 peptides undergo structural conformation changes when mixed

In order to investigate the conformations of the peptides in the various systems, we used a two-stage machine learning protocol (UMAP (29) and HDB-SCAN (30)) to cluster the conformations taken by each peptide over the course of each simulation. The resulting conformations were further characterised using the DSSP algorithm for secondary structure prediction (35). It is important to note that DSSP is unable to identify PII conformations, which is a distinctive feature of WF AMPs (13).

Clustering of WF1a conformations in both pure and mixed systems revealed 12 distinct conformation clusters without any outliers (Fig. SI 2 a, b, c). All but one of these were found in the pure systems, with conformations 7(29.83%), 10(9.41%) and 12(28.44%) making up most of the structures. The conformations observed are predominantly unstructured, with certain clusters containing an *α*-helical region at either the N-terminus (conformation 12) or C-terminus (conformation 10) (Fig. 2 a).

**Fig. 2.**
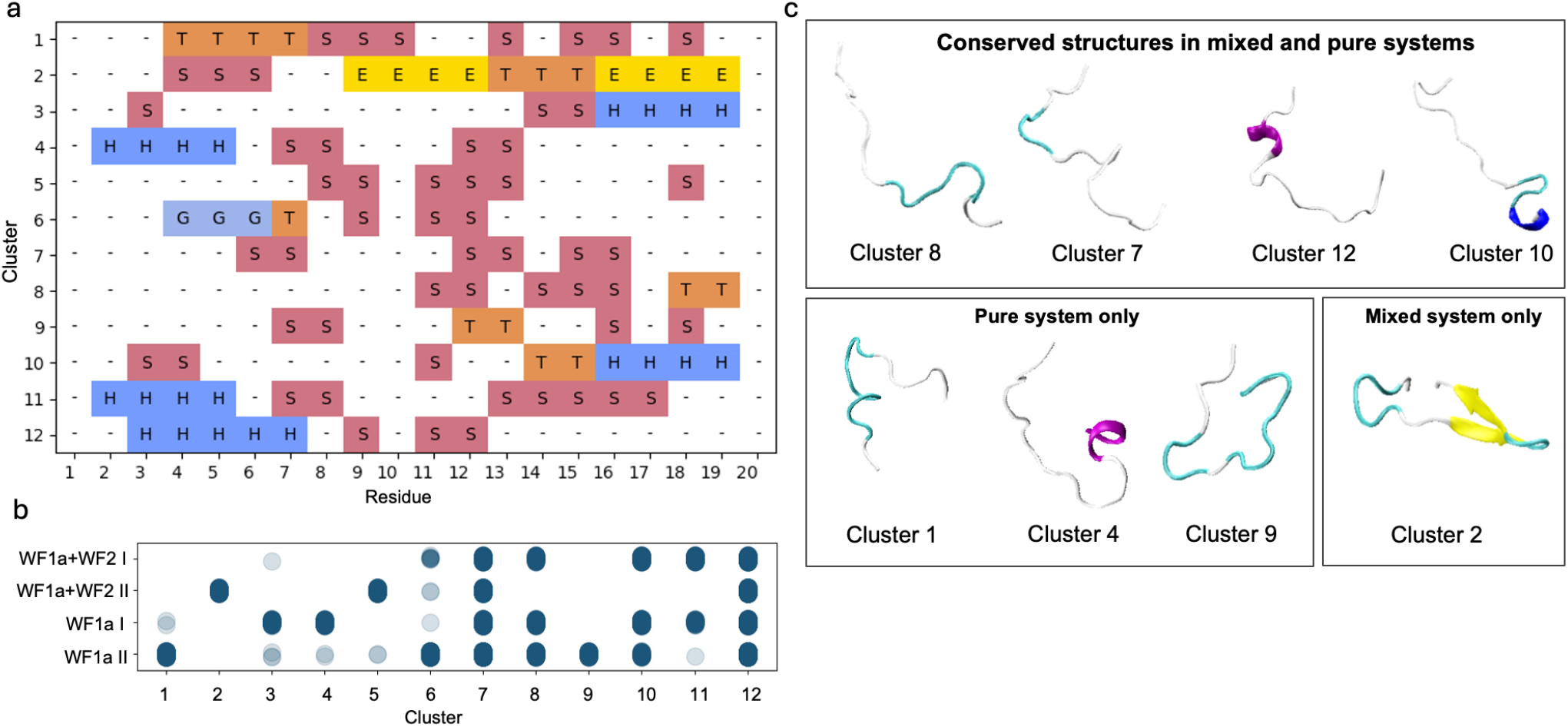
Conformational clusters observed for WF1a peptides. (a) Representative secondary structures (b) The distribution of conformations per system and simulation, and the secondary structure assignment per residue for each cluster. The meaning of each letter can be found in Table 2.

Combining the two AMPs did not greatly alter the conformations adopted by the WF1a peptides, with most conformations present in both type of systems (Fig. 2 b). Exceptions were conformation 2, containing a *β*-sheet structure, which is exclusive to the mixed systems, and conformations 1, 4 and 9, which were present only in the pure system. Only conformations 4, 11 and 12 adopt a short *α*-helical conformation at their N-terminus, while two conformations, namely conformation 3 and 10 adopt an *α*-helical conformation at their C-terminus. The rest of their structures display mainly disordered regions or bends and turns. No *α*-helices were noted in the central region of the peptide. In contrast, the structural conformations of WF2 peptides were spread into a larger number of smaller clusters ranging from 13% of the conformations, to as low as 2%, such as conformation 4 (Fig. SI 2 d, e, f). WF2 peptides adopt more structures in the pure systems compared to WF1a peptides. Half of the conformations observed in the pure WF2 systems contain a substantial amount of *α*-helical content, while the rest of the conformations are largely unstructured.

Mixing the two types of peptides has a stronger effect on the conformations adopted by the WF2 peptides, as evidenced by a limited overlap in conformations observed between the mixed and pure systems (Fig. 3 b). The common conformations include conformations 14, 15, 17 and 20. Conformations 14 and 15 are disordered while conformations 17 and 20 have a large *α*-helical content. Some of the structures adopted by WF2 peptides in the mixed systems seem to lose part of their helical content, such as conformations 1, 4, 5, and 11. Overall, the loss of *α*-helical content does not appear to be specific to either type of system.

**Fig. 3.**
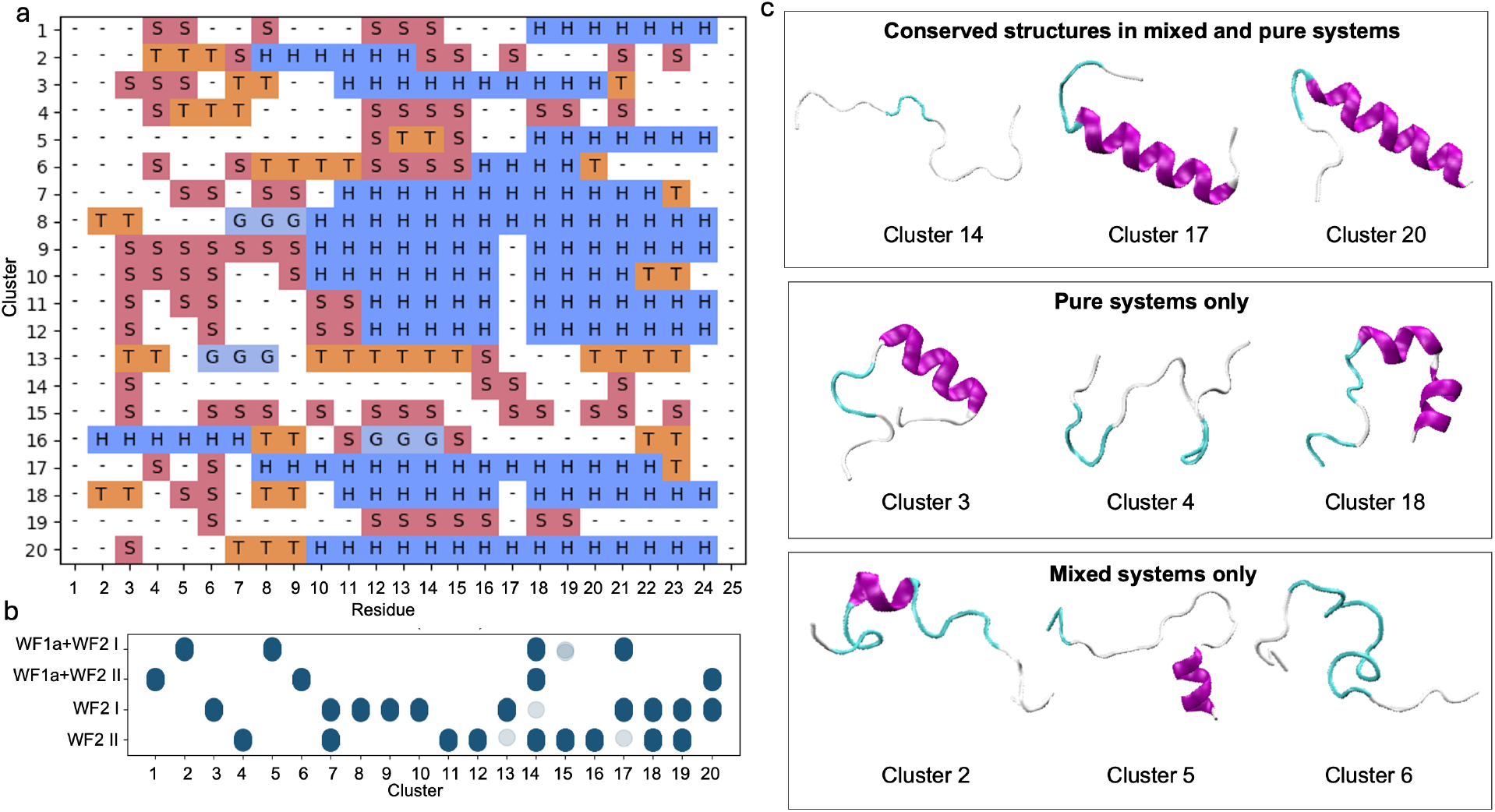
Conformational clusters observed for WF2 peptides. (a) Representative secondary structures (b) The distribution of conformations per system and simulation, and the secondary structure assignment per residue for each cluster. The meaning of each letter can be found in Table 2.

### F. Conformation affects the orientation of the peptides relative to the membrane

Both peptides are able to insert into the membrane, with deeper insertion seen in WF2 peptides in both the pure and mixed systems (Fig. 4 a). However, we found that the orientation of the peptides relative to the lipid bilayer is dependent not only on whether the peptides are acting alone or with their synergistic pair, but also the conformation that the peptide takes during insertion (Fig. 4 b).

**Fig. 4.**
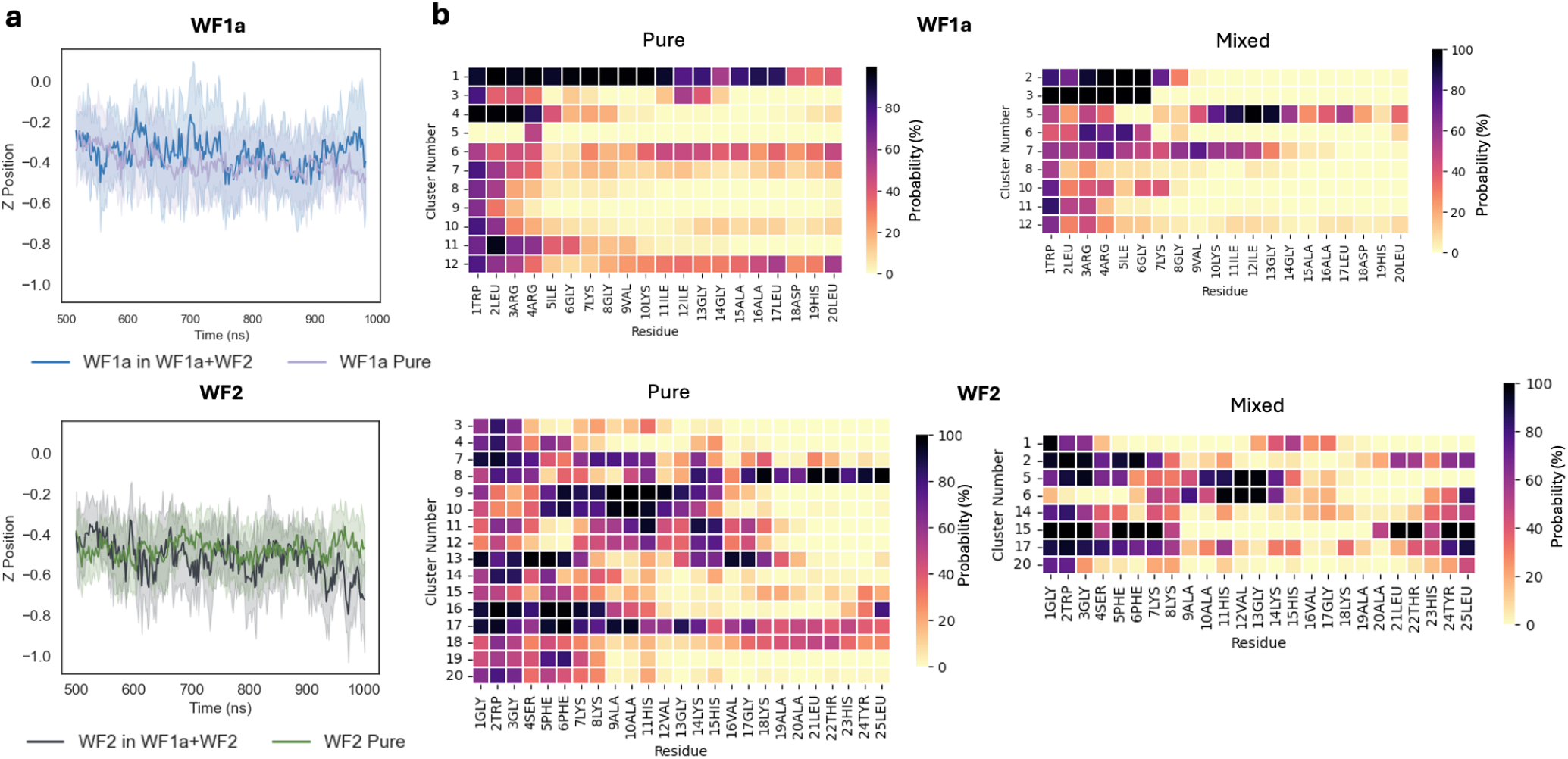
(a) The depth of insertion into each membrane is shown as the Z-position over time. Positive or negative values indicate the peptides are below or above the phosphate group. (b) Probability of inserting below the the phosphate group of each residue in each conformation.

In the pure systems WF2 peptides are more likely to insert below the phosphate plane of the membrane lipids via their N-terminus residues. However, we noted a number or conformations which are more likely to insert via the middle residues (conformational clusters 9, 10, 11, 12, 13 & 17).

Other conformations are able to penetrate the bilayer via either termini, but this behaviour is restricted to fewer conformational clusters (8, 16, 17 & 18). Mixing the two peptides causes WF2 to change orientation and insert into the membrane predominantly via either termini, although some variation between the different conformations still exists. WF1a conformations displayed the same heterogeneity in their orientation while simulated individually. Three of the WF1a conformations are able to insert into the membrane with most of their residues (conformational clusters 1, 6 & 11), while the rest of the conformations insert either via the N-terminus or a combination of N- and C-terminus. Mixing the peptides leads to a shift towards insertion almost exclusively via the N-terminus. Only conformations 5 and 7 are likely to insert the membrane via a larger proportion of their residues.

In addition to our analysis of membrane insertion, we investigated the effects of mixing the two peptides on the membrane. We conducted analyses of the hydrogen bonding between the peptides and the bilayer lipids (Fig. SI 3 a, c), the area per lipid of the membrane (Fig. SI 3 b) and the membrane curvature (Fig. SI 4). We only observed minimal changes between the different systems at the time scale studied.

### G. The interplay between aggregation and conformation in WF1a and WF2 peptides

We investigated the relationship between conformations and aggregation propensity, which unveiled that only a limited subset of conformations exclusively existed in the monomeric state (Fig. 5). Specifically, in the case of WF1a peptides, we found that conformation 5, a largely unstructured conformation is the least likely conformation to be involved in aggregates. All of the other conformations seem to be found in aggregates of different sizes, in various proportions. Therefore, there does not seem to be any straightforward relationship between AMPs structure and their ability to aggregate.

**Fig. 5.**
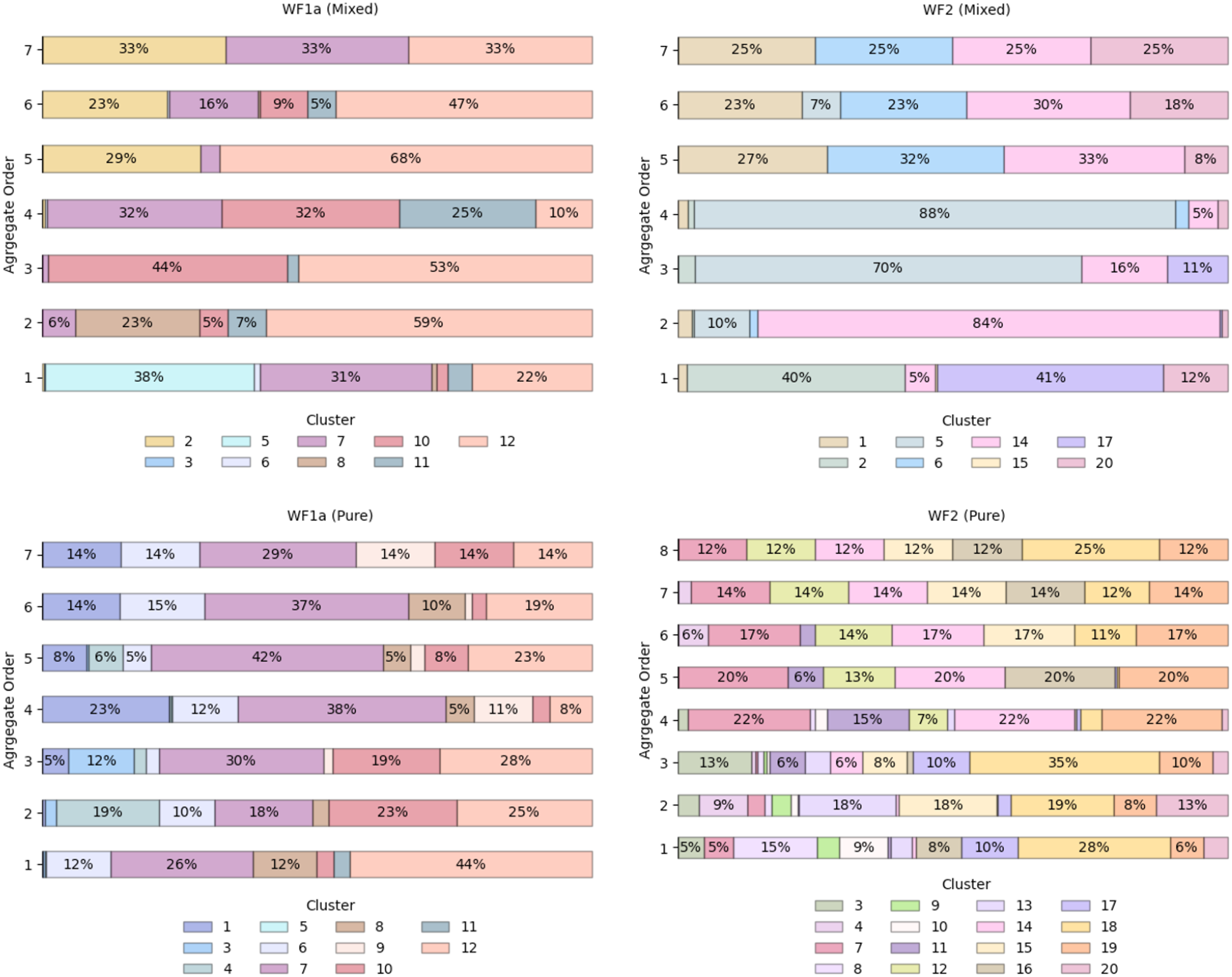
The distribution of WF1a and WF2 conformations based on the size of the aggregates.

In the case of WF2 peptides, only conformation 17 was observed to have a preference for monomers and lower-order aggregates. Even though WF2 peptides are likely to adopt an *α*-helical structure, a number of other conformations that are mostly unstructured are also able to aggregate. Therefore, the presence of *α*-helical structure or the lack thereof does not dictate the ability of WF2 to aggregate.

Further investigation into the distribution of conformations across different aggregate orders showed a preference among certain conformations for higher-order aggregates, while several conformations were present across various aggregate orders (Fig. 5a). For instance, in the mixed systems, WF1a conformation 2 was exclusively present in higher-order aggregates such as 5-mers, 6-mers, and 7-mers. Similarly, WF2 conformations 1 and 6 are predominantly associated with aggregates of order *n >* 4. Conversely, several conformations exhibited the ability to exist as both monomers and oligomers, as shown by WF1a conformations 7, 10, 11 & 12, and WF2 conformations 2, 12, 14 & 20.

Specific WF2 conformations are centrally located within aggregates, interacting with more than two peptides simultaneously (Table SI 1). Two WF2 conformations, namely conformation 12 in the pure WF2 system, and conformation 14 in the mixed system, appear to be central to the aggregates in which they are involved. Conformation 14 exhibits a highly disordered secondary structure, while conformation 12 adopts a predominantly *α*-helical structure, with bends and disordered regions at the N-terminus, and an *α*-helical conformation at the C-terminus, suggesting that a disordered structure is not a requirement for the WF2 peptides to be able to interact with multiple peptides at the same time. Further, we also observed that, in higher-order heteromers such as 7-mers (Fig. 6), the peptides interact preferentially with peptides of the same type, resulting in the formation of two sub units linked by a specific WF1a-WF2 pair. Hence, specific conformation pairs act as a linker and play a crucial role in facilitating the formation of higher-order aggregates.

**Fig. 6.**
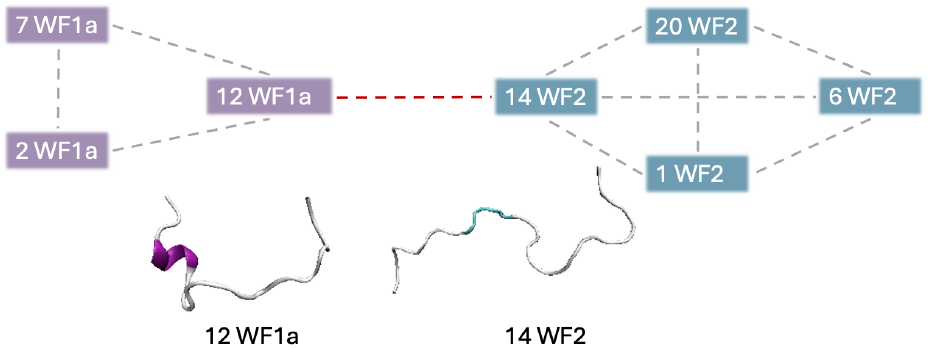
Graphical representation of the interaction pattern between the conformations observed in 7-mer heteromers.

### D. Combining WF1a and WF2 leads to a greater number of intermolecular interactions in heteromers

Certain combinations of conformations are more prone to interactions, and in particular, WF1a conformations are more likely to be be found in contact with other WF1a conformations (Table SI 2) than with WF2 peptides. WF1a peptides with conformation 4 are seen to bind specifically with conformation 10 WF1a peptides, and conformation 7 are found to bind with conformation 9 peptides in the pure system. In the mixed system WF1a conformation 2 is found to specifically bind with conformation 12 and conformation 7 is found to bind with conformation 7. These are the most commonly found interacting pairs of conformations for all of the pairs observed in our simulations.

The two AMPs have different modes of interaction with the other peptides. In mixed systems a larger number of residues are involved in the WF1a-WF1a interactions than are found in the WF1a-WF2 and WF2-WF2 interactions, particularly notable in the C-terminus region (Fig. SI 4). This trend persists in the pure systems, where WF1a-WF1a interactions occur via a larger proportion of residues than WF2-WF2 interactions (Fig. SI 5). Therefore, WF1a peptides are able to interact with other peptides via more residues, which might allow for stronger and more stable intermolecular interactions than those involving only WF2 peptides.

The peptide interaction analysis shows distinctive interaction preferences influenced by the system composition and peptide type (Fig. 7). WF2 peptides tend to interact with other WF2 peptides via both the N- and C-terminus, while the interactions with WF1a peptides occur mostly using their C-terminus residues. In contrast, WF1a peptides interact with other peptides predominantly via the help of the residues nearest the C-terminus. Interestingly, in the mixed systems WF1a peptides interact via different amino acids depending on whether it is interacting with another WF1a peptide or a WF2 peptide. Interactions of WF1a peptides with WF2 peptides predominantly involve residues 9VAL, 10LYS, and 12ILE, while interactions with analogous peptides primarily include the hydrophobic residues 14GLY, 15ALA & 16ALA.

**Fig. 7.**
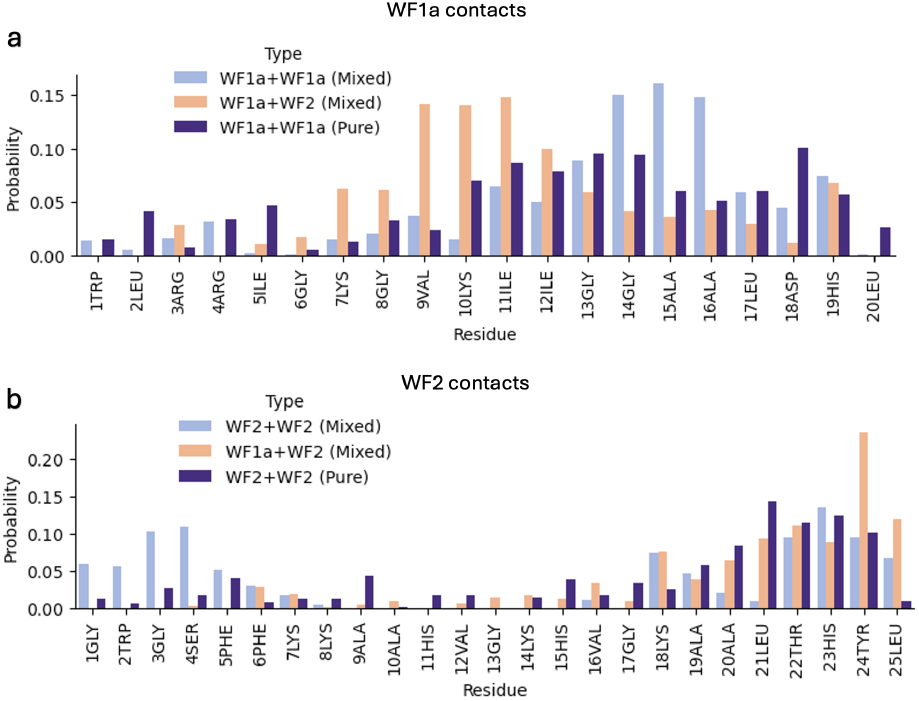
The probability of being involved in contacts per each amino acid residue for the two AMPs. The probability of WF1a residues contacts in WF1a-WF1a, WF1a-WF2 in the pure and mixed systems (a). The probability of WF2 residues contacts in WF2-WF2 and WF1a-WF2 in the pure and mixed systems (b).

Mixing the two peptides leads to a greater number of interactions (Fig 8). WF2 peptides have a greater ability to form hydrogen bonds and electrostatic interactions when bound to WF1a peptides or other WF2 peptides. In contrast, in the mixed system, WF1a-WF1a interactions are mostly driven by the hydrophobic effect. Interestingly, no hydrogen bonding capabilities were observed in the WF1a-WF1a and WF2-WF2 interactions in the pure system. This suggests that combining the two peptides results in stronger and more stable oligomers.

**Fig. 8.**
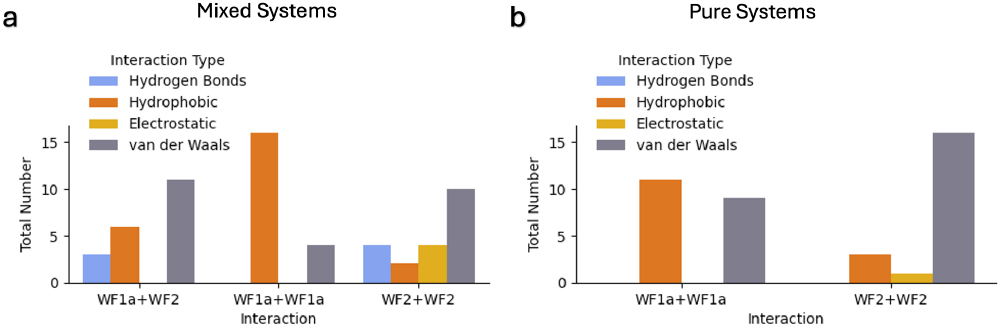
The distribution of the different types of interactions in the mixed (a) and pure (b) systems.

The analysis of the mixed systems indicates that interactions between certain conformations primarily arise from the hydrophobic effect, while others involve a combination of hydrogen bonding and hydrophobic interactions. Some conformation pairs, like WF1a cluster 8 with WF2 conformation 14 and WF1a conformation 11 with WF2 conformation 5, engage mostly through hydrogen bonding, with 3ARG:24TYR and 10LYS:22THR being amongst the most probable interactions. The interaction between WF1a and WF2 peptides often relies on one or two hydrophobic contacts, which draw together surrounding residues to form weaker van der Waals interactions. These results suggest that the conformations of the peptides play a crucial role in the type of interactions the two AMPs engage in.

In terms of interactions of peptides of the same type, WF2-WF2 contacts are influenced by the conformation of the interacting peptides (Table SI 5 and Table SI 6). For example, interactions between conformations 1 and 20 involve a hydrogen bond at 4SER:22THR and a hydrophobic interaction at 6PHE:19ALA. Other conformations may interact through electrostatic interactions, including *π − π* stacking, driven by residues such as 4SER, 5PHE, 7LYS, 18LYS, 23HIS, and 24TYR. In pure systems, most conformation interactions occur via van der Waals forces. In contrast, the interactions between WF1a-WF1a peptides are predominantly driven by hydrophobic effect, both in pure and mixed systems, and are not significantly influenced by the specific conformations of the WF1a peptides (Table SI 3 and Table SI 4).

## Discussion

This study investigated the interplay between the aggregation behaviour and structural dynamics of the synergistic pair of WF1a and WF2 AMPs, shedding light on the potential mechanisms of their interactions. AMPs are crucial components of the innate immune system, that have gained attention for their ability to combat microbial infections (39–41). In this context, understanding the mechanisms underlying AMP activity, including peptide conformation and aggregation, and their synergistic effects, holds promise for developing novel antimicrobial strategies.

The formation of aggregates has been proposed as key in the antimicrobial activity of AMPs and possibly to the mechanism of synergy between AMPs (42). Our findings reveal that mixing the two peptides induces a shift towards the formations of higher-order aggregates. When mixed, WF1a peptides demonstrate a greater probability aggregating than WF2, albeit typically in lower-order oligomers. Conversely, WF2 peptides exhibit a lower probability of association but when they are involved in aggregates, they have a preference for forming higher-order aggregates. This suggests that WF1a peptides help WF2 peptides create larger aggregates, a behaviour previously reported in the case of PGLa and MAG2, where the presence of MAG2 might increase the aggregation of PGLa peptides in a mixture (43).

Building upon this observation, we noted a preference for heteromer formation over homomer assembly, a phenomenon previously documented in other AMP systems such as MAG2 and PGLa(44, 45). Even though MAG2 and PGLa can form pores on their own and have antimicrobial activity, their synergistic activity leads to the formation of a stronger heteromer, which has been linked to stronger antibacterial activity (46). Therefore, we expect that the formation of heteromers between WF1a and WF2 peptides is an important step in their synergistic activity. Interestingly, we also noted that the ratio of WF1a:WF2 peptides in the formed heteromers changes depending on the aggregate size, with higher-order aggregates containing a higher proportion of WF2 peptides than smaller oligomers.

The secondary structure of AMPs play an important role in their antimicrobial potency (47). Certain AMPs undergo conformational changes upon interacting with lipid membranes, transitioning from disordered to helical conformations, which is crucial for their activity. For instance, mastoparans from wasp venom and GL13K from human parotid secretory protein adopt *α*-helical structures upon membrane binding, emphasizing the significance of secondary structure in mediating antimicrobial activity (48, 49).

Our analysis of the structural dynamics of the two WF AMPs reveals that mixing the two peptides has a strong effect on their conformations, particularly for WF2 peptides. Mixing the two peptides constrains the conformational space of WF2 peptides, which adopts a lower number of conformations than when simulated alone. WF2 peptides are predominantly characterized by *α*-helical structures and have been shown to exhibit substantial conformational flexibility (13), a distinctive feature relative to MAG2 (18) or temporins (42). WF2 peptides undergo more conformational transitions than WF1a peptides when simulated individually, but only undergo limited conformational transitions in the mixed systems. WF2 peptides display a higher tendency to form *α*-helical structures, which is limited at the N-terminus. This may be attributed to the N-terminus involvement in membrane insertion, a known characteristic of WF2 peptides (13, 18), which we also observed in our analyses (Fig. 4 b). In contrast, WF1a peptides predominantly display disordered conformations, with sporadic helical regions at either termini. Interestingly, one WF1a conformational cluster observed in the mixed systemz adopted a *β*-sheet structure, which has been observed in multiple AMPs, such as 1-purothionin (50) and gomesin (51). Mixing the peptides constrains their conformational space, reducing the number of distinct conformations adopted by the peptides.

Further, the conformation of the peptides and whether the peptide is acting alone or with its synergistic pair also affects the orientation of the peptides and their insertion into the membrane. When mixed, WF2 peptides gain an ability to insert via both termini, while WF1a peptides lose their ability to insert via the C-terminus, and predominantly insert via the N-terminus. It is also possible that the orientation of the peptides is further influenced by their aggregation behaviour. The ability of AMPs to aggregate can be affected by modifications of their sequence that alter their structure, such as N-terminus lipidations (52) and carboxylation and conversion of C-terminus cysteine (53). This suggests that the structures adopted by AMPs are essential to their ability to aggregate. However, in our case, no clear link between specific peptide structures and ability to aggregate was observed in the WF1a and WF2 peptides. The predominant conformations observed in higher-order aggregates differs from those in monomers for both WF1a and WF2 peptides. Additionally, our investigation suggests the potential formation of higher-order aggregates through the combination of two sub-aggregates composed solely of peptides of one type, interconnected by a linker formed between one WF1a peptide and one WF2 peptide.

Further, WF2 peptides are engaged in stronger intermolecular interactions, such as hydrogen bonds with both WF1a and WF2 peptides, while WF1a peptides primarily interact with other WF1a peptides via hydrophobic and weaker van der Waals interactions. Electrostatic interactions such as salt bridges between anionic and cationic residues in membrane-bound peptides, and proteins have been shown to play a role in the assembly of oligomers (43, 54–56). Therefore, it appears that the involvement of WF2 peptides is important for establishing stronger intermolecular contacts within aggregates. We also noted that certain WF2 conformations exhibit a preference for specific intermolecular interactions, indicating their ability to undergo conformational changes to accommodate specific peptide interactions.

Interestingly, WF1a peptides are able to interact via a larger proportion of their residues with either WF1a or WF2 peptides, possibly leading to more stable interactions. WF1a interacts with other peptides mainly by its middle and C-terminus residues, likely facilitated by the extended conformations it adopts. WF1a peptides may protect WF2 peptides by adopting an unfolded state and wrapping them using a larger number of residues, akin to the mechanism proposed for distinctin, a heterodimer linked by a disulfide bridge (57). One of distinctin’s two chains adopts an extended conformation that protects the dimer from proteolytic digestion (58). Despite the contribution with stronger intermolecular interactions of WF2 peptides, WF1a’s broader structural involvement underscores its importance in heteromer stability.

Despite the prevalent formation of higher-order aggregates, a number of WF2 peptides remain in a monomeric state. These monomers may have the ability to penetrate the bacterial membrane more deeply compared to aggregates, while aggregates could contribute to the destabilization of the bacterial membrane (Fig. 4 a). Although AMPs aggregates have been shown to be important for the peptides antibacterial activity by destabilizing the bacterial membrane in certain cases, a number of studies have shown that self-association or aggregation of AMPS might compromise the peptides efficacy, possibly due to a reduced ability to translocate the bacterial membrane (59–61).

However, complementing our findings with the effects caused by the peptides on the membrane lipids would further deepen our understanding of how synergistic AMPs work. Apart from an increase of penetration into the membrane by WF2 peptides when the peptides are mixed and the changes in orientation of both peptides, our investigation of the effects on the membrane lipids showed only modest changes, not allowing us to make any definite links between the peptides interactions, their conformations and their effects on the membrane. One possible reason could be the time scale studied of just 1 microsecond, which might not be long enough to allow the peptides to exert any significant effects on the membrane. For example, a number of AMPs are known to form membrane pores, resulting in the loss of membrane potential and rapid release of intracellular components and death (62–64). The timescale of pore formation ranges from microseconds to seconds, much longer time frames than presented in our work.

Increasing the duration of the simulations might allow the peptides to further insert into the membrane and more pronounced membrane disruption to happen. The analysis of AMPs aggregation and conformational flexibility could then be complemented with their impact on bacterial membrane, to provide further insights into their mechanisms of action. Further, the structure and interactions of AMPS can also be affected by their environment, including the membrane lipid structure and packing (65). Moreover, it has been suggested that the formation of higher-order aggregates or ‘supramolecule’ arrangements is strongly dependent on the membrane composition (66). Therefore, the study of the conformation dynamics and aggregation behaviour of synergistic AMPs in other models of bacterial membranes might provide more insights into their behaviour.

## Conclusion

In summary, our study provides a computational method for isolating and characterising the conformations of AMPs using a combination of molecular dynamics simulations and unsupervised machine learning. We provide new insights into the aggregation behaviour and the structural dynamics of WF1a and WF2 antimicrobial peptides that can help understand their synergistic interactions. We highlight the significance of heteromer formation, peptide conformational diversity, and their implications for antimicrobial strategies. Further exploration of AMP aggregation and structure dynamics and their impact on bacterial membranes is crucial for advancing our understanding and developing effective antimicrobial agents.

## Author contributions

Miruna Serian: Data curation, formal analysis, investigation, methodology, software, validation, visualisation, writing - original draft. A. James Mason: Conceptualization, supervision, writing - review & editing. Christian D. Lorenz: Conceptualization, funding acquisition, project administration, resources, supervision, writing - review & editing.

## Data availability

Simulations files and analyses scripts are available at [this repository](https://github.com/Lorenz-Lab-KCL/publications/tree/main/AMPSynergy2024_manuscript_rsc).

## Acknowledgements

We are grateful to the UK High-End Computing Consortium for Biomolecular Simulation (HEC BioSim), which is funded by EPSRC (EP/R029407/1), and UK Materials and Molecular Modelling Hub for computational resources, which is partially funded by EPSRC (EP/T022213/1, EP/W032260/1 and EP/P020194/1) for providing us access to computational resources. This work also benefited from access to the King’s Computational Research, Engineering and Technology Environment (CREATE) at King’s College London (67). This work was supported by the Biotechnology and Biological Sciences Research Council (BB/T008709/1) via the London Interdisciplinary Doctoral Programme (LIDo). Supplementary information

**ACKNOWLEDGEMENTS**

