## Supplemental Data for "Unsupervised learning elucidates the interplay between conformational flexibility and aggregation in synergistic antimicrobial peptides"

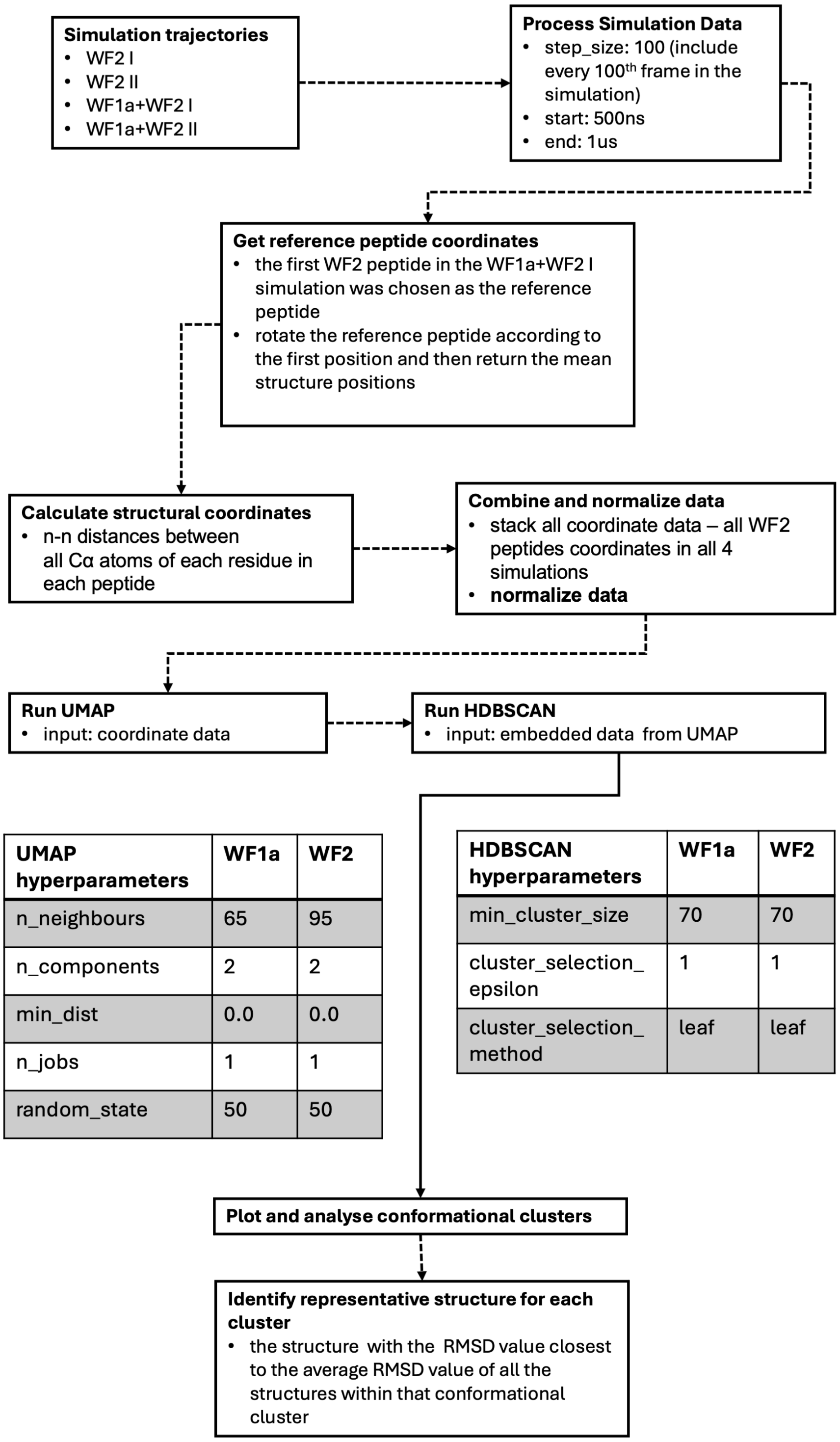


| UMAP hyperparameters | WF1a | WF2 |
|---|---|---|
| n_neighbours | 65 | 95 |
| n_components | 2 | 2 |
| min_dist | 0.0 | 0.0 |
| n_jobs | 1 | 1 |
| random_state | 50 | 50 |

| HDBSCAN hyperparameters | WF1a | WF2 |
|---|---|---|
| min_cluster_size | 70 | 70 |
| cluster_selection_epsilon | 1 | 1 |
| cluster_selection_method | leaf | leaf |

**Fig. SI 1**: Example of UMAP and HDBSCAN workflow for WF2 peptides. A similar workflow was applied for WF1a peptides.

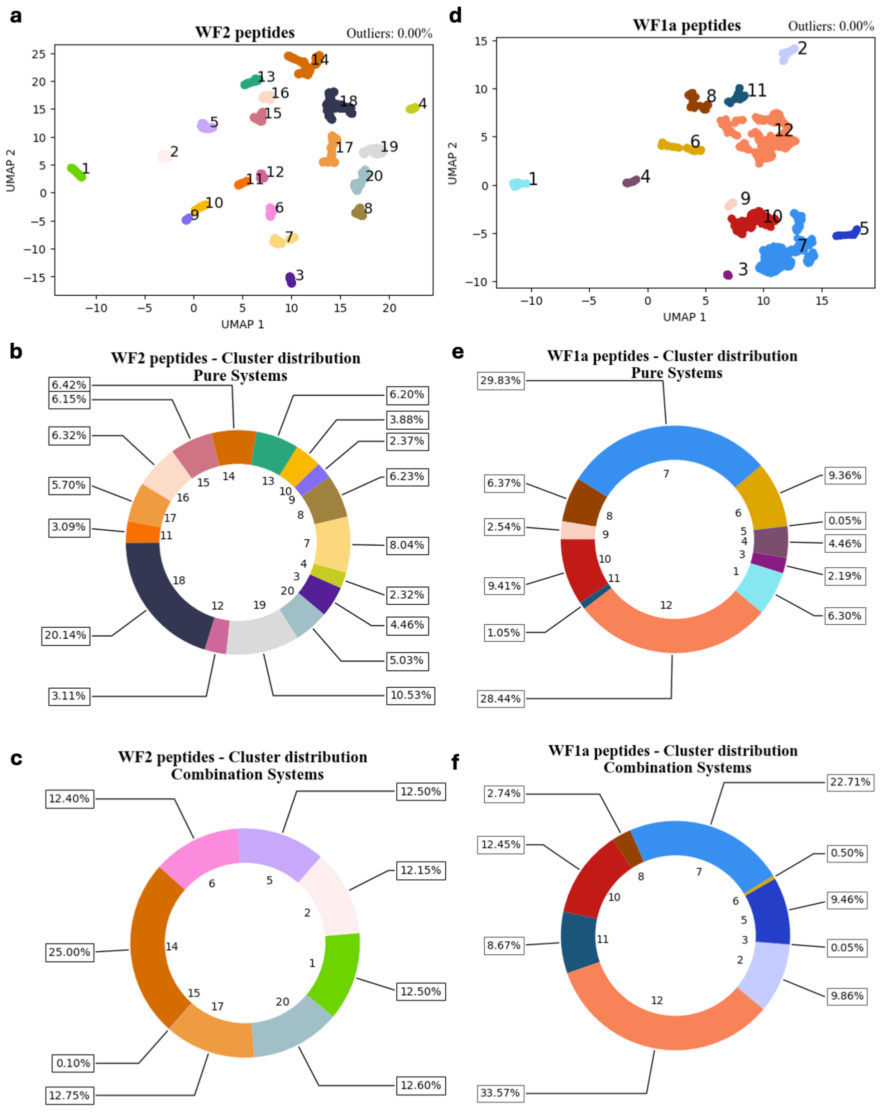


**Fig. SI 2**: Conformational clusters. UMAP projections in 2 dimensions for WF2 (a) and WF1a (b). The cluster distribution as percentages for WF2 - Pure systems (b), WF2 - Combination Systems (c) and WF1a - Pure Systems (e), WF1a - Combination Systems (f).

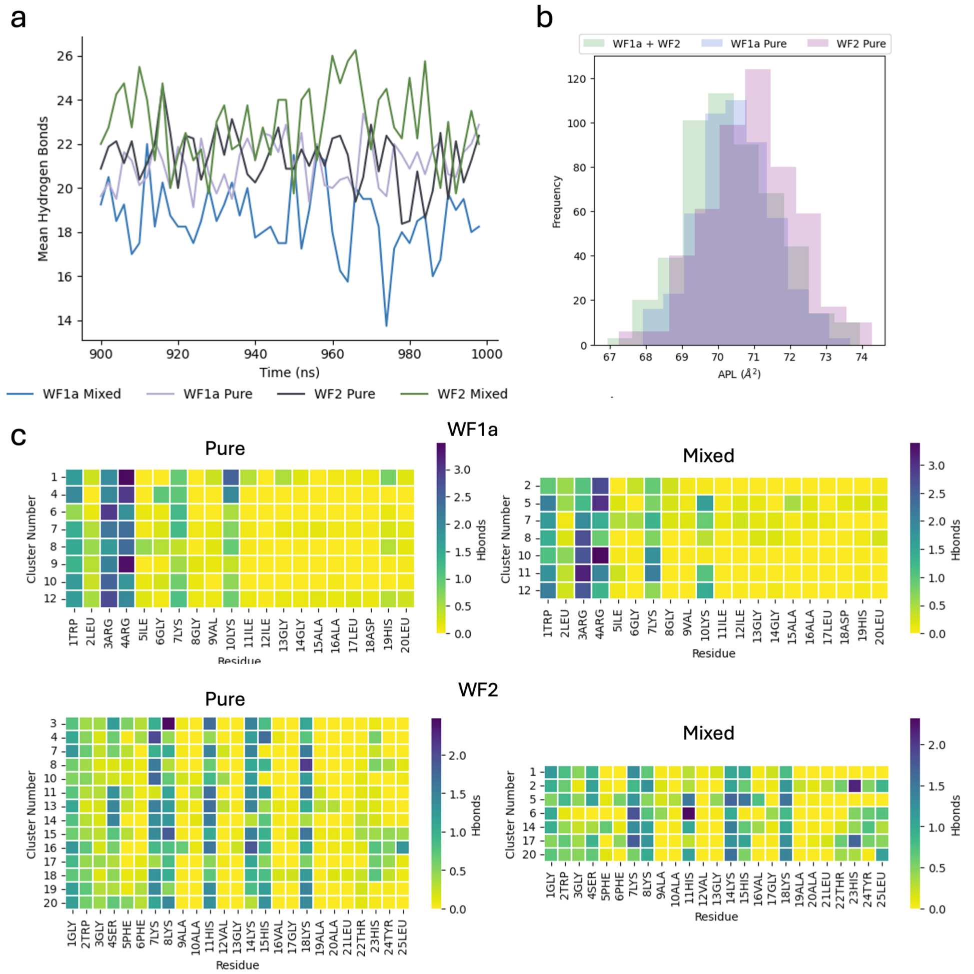


**Fig. SI 3**: Mean number of hydrogen bonds made with the membrane lipids per type of peptide and system during the last 100 ns of the simulations (a). Area per lipid (APL) distribution for each system during the last 50 ns of the simulations (b). Mean hydrogen bonds with membrane lipids per each residue and conformation (c).

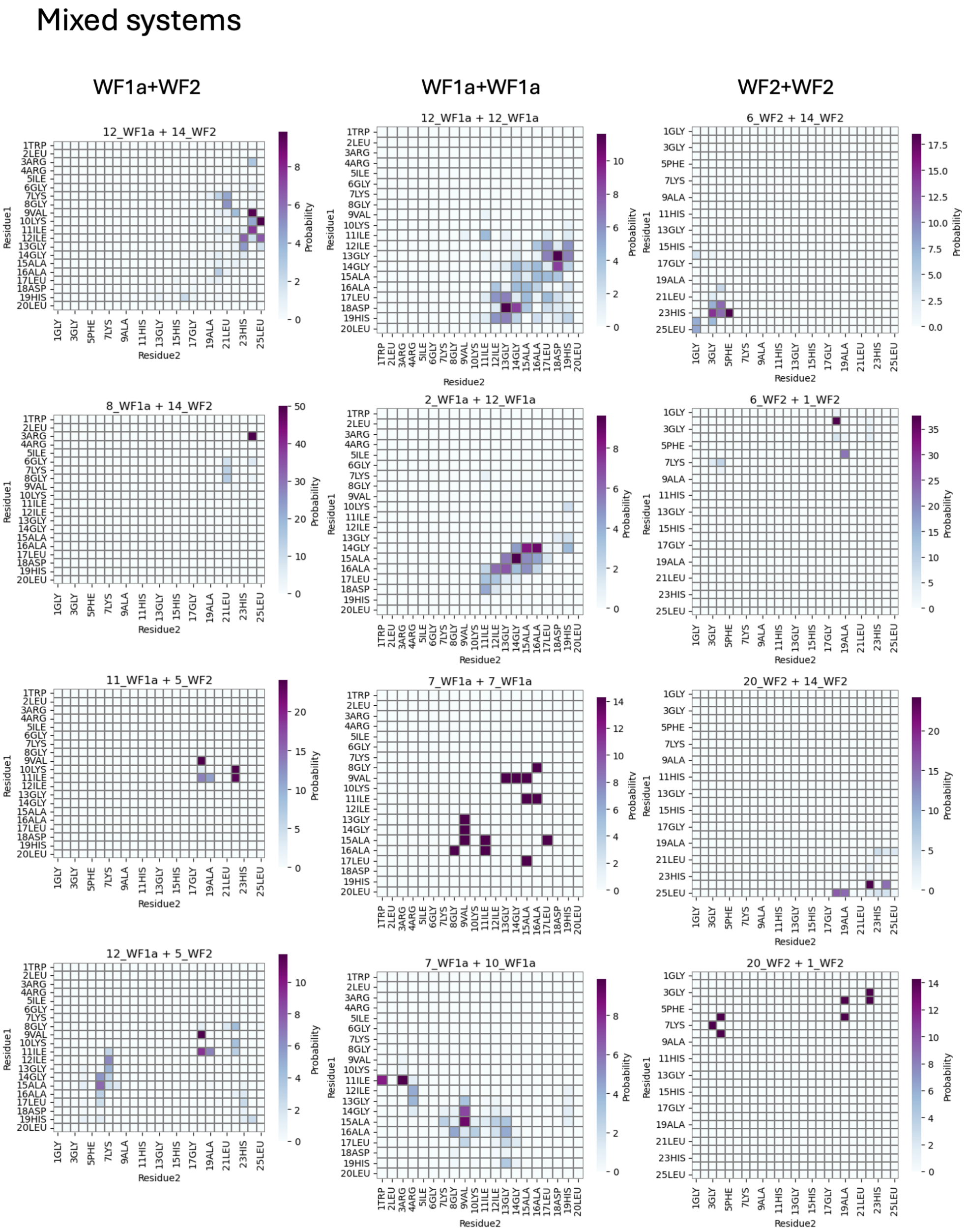


**Fig. SI 4**: Cluster-Cluster interactions in the mixed systems.

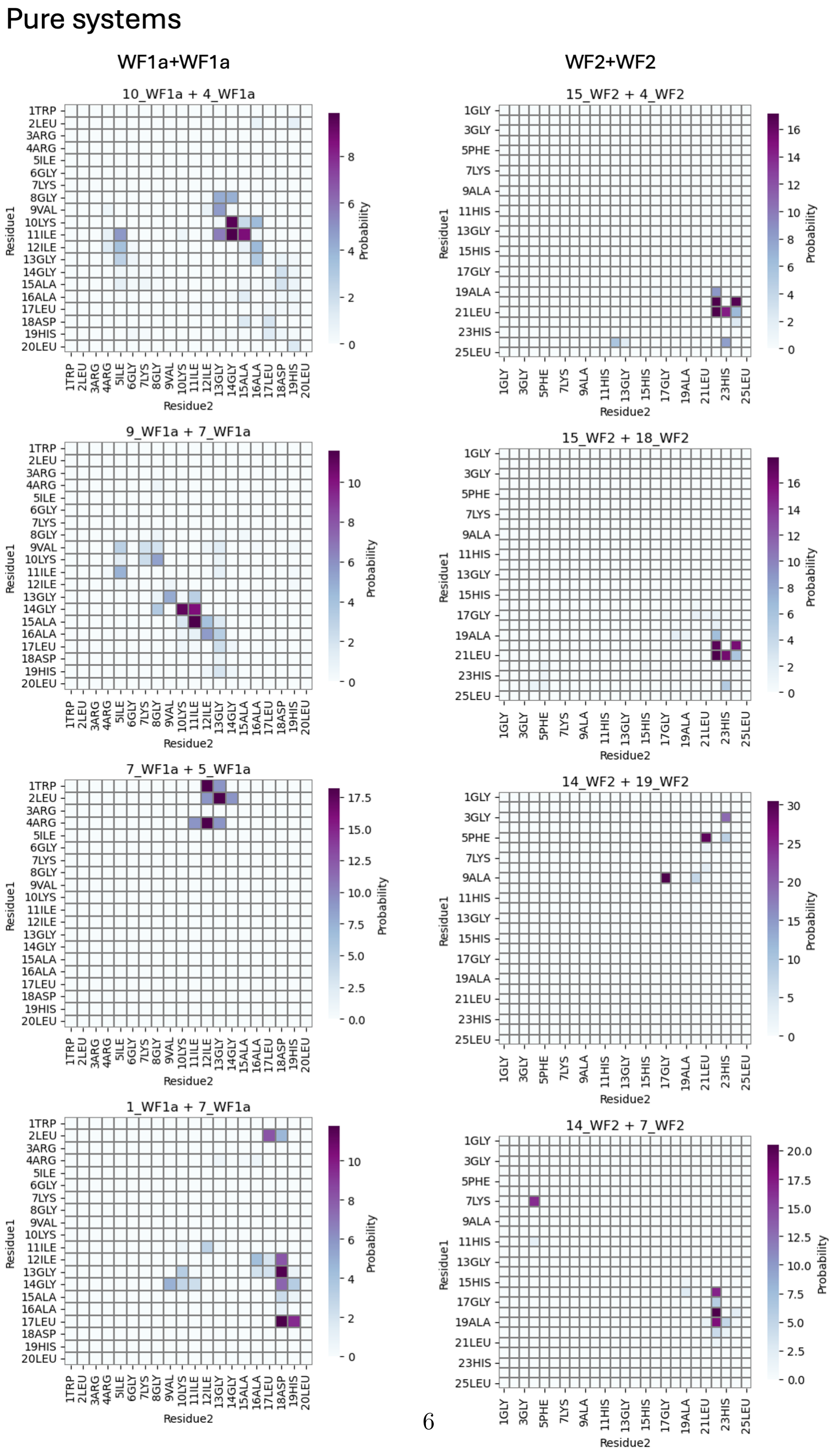


**Fig. SI 5**: Cluster-cluster interactions in the pure systems grouped by the types of peptides involved.

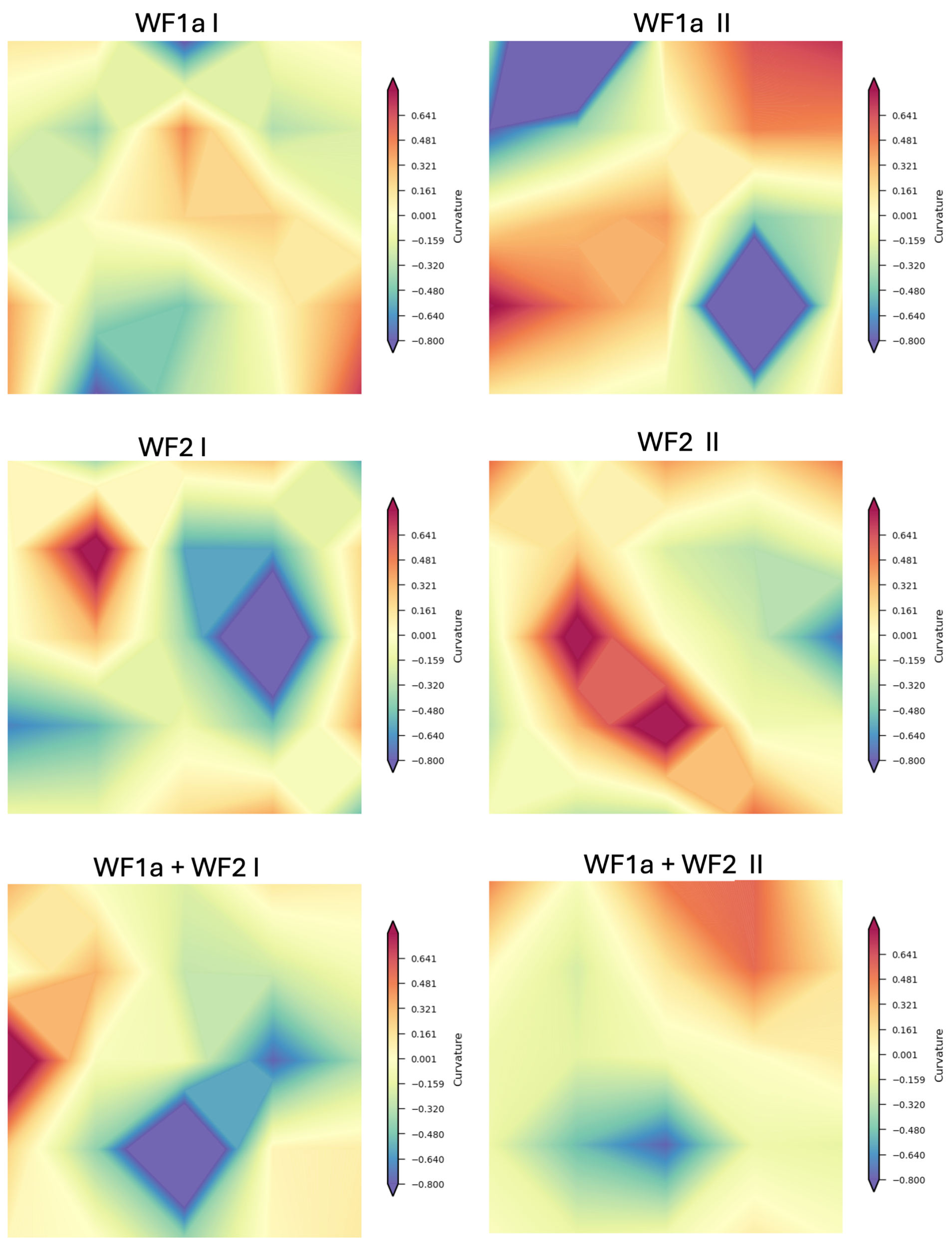


**Fig. SI 6**: The membrane curvature of the upper leaflet at 1 microsecond for each system and simulation. Negative values indicate a negative curvature while positive values indicate a positive curvature. The curvature data was generated using the Lipid-Dyn Package [1]

| **Cluster** | **WF1a** **Combination** **Mean** | **SD** | **Pure** **Mean** | **SD** |
|---|---|---|---|---|
| **1** | | | 1.69 | 0.65 |
| **2** | 1.77 | 0.66 | | |
| **5** | 1.79 | 0.60 | 1.0 | 0.0 |
| **6** | 1.33 | 0.47 | 1.81 | 0.70 |
| **7** | 1.53 | 0.53 | 1.99 | 0.74 |
| **8** | | | 1.14 | 0.35 |
| **9** | | | 2.04 | 0.94 |
| **10** | | | 1.46 | 0.50 |
| **12** | 2.13 | 0.54 | 1.32 | 0.49 |

(a) WF1a peptides

| **Cluster** | **WF2** **Combination** **Mean** | **SD** | **Pure** **Mean** | **SD** |
|---|---|---|---|---|
| **1** | 1.51 | 0.69 | | |
| **4** | | | 1.00 | 0.00 |
| **6** | 1.82 | 0.38 | | |
| **7** | | | 1.9 | 0.3 |
| **11** | | | 1.93 | 0.59 |
| **12** | | | 2.67 | 0.49 |
| **14** | 3.17 | 0.63 | 2.87 | 0.50 |
| **15** | | | 1.94 | 0.37 |
| **16** | | | 1.11 | 0.31 |
| **17** | | | 1.00 | 0.0 |
| **18** | | | 1.16 | 0.47 |
| **19** | | | 1.23 | 0.42 |
| **20** | 1.36 | 0.48 | | |

(b) WF2 peptides

**Table SI 1**: Cluster centrality values for WF1a (a) and WF2 peptides (b). The values were computed using the degree_centrality() function of NetworkX library [2]

| Cluster_pep1 | Cluster_pep2 | Probability % | System | pep1 | pep2 |
|---|---|---|---|---|---|
| 12 | 14 | 5.12 | WF1a_WF2 | WF1a | WF2 |
| 11 | 5 | 3.27 | WF1a_WF2 | WF1a | WF2 |
| 12 | 5 | 2.19 | WF1a_WF2 | WF1a | WF2 |
| 8 | 14 | 1.67 | WF1a_WF2 | WF1a | WF2 |
| 2 | 12 | 8.04 | WF1a_WF2 | WF1a | WF1a |
| 7 | 7 | 7.00 | WF1a_WF2 | WF1a | WF1a |
| 7 | 10 | 4.38 | WF1a_WF2 | WF1a | WF1a |
| 12 | 12 | 2.30 | WF1a_WF2 | WF1a | WF1a |
| 12 | 14 | 5.12 | WF1a_WF2 | WF1a | WF2 |
| 11 | 5 | 3.27 | WF1a_WF2 | WF1a | WF2 |
| 12 | 5 | 2.19 | WF1a_WF2 | WF1a | WF2 |
| 8 | 14 | 1.67 | WF1a_WF2 | WF1a | WF2 |
| 10 | 4 | 9.55 | WF1a_only | WF1a | WF1a |
| 9 | 7 | 7.88 | WF1a_only | WF1a | WF1a |
| 7 | 5 | 5.50 | WF1a_only | WF1a | WF1a |
| 1 | 7 | 4.68 | WF1a_only | WF1a | WF1a |
| 6 | 14 | 4.04 | WF1a_WF2 | WF2 | WF2 |
| 6 | 1 | 1.87 | WF1a_WF2 | WF2 | WF2 |
| 20 | 14 | 1.20 | WF1a_WF2 | WF2 | WF2 |
| 20 | 1 | 0.03 | WF1a_WF2 | WF2 | WF2 |
| 15 | 4 | 5.84 | WF2_only | WF2 | WF2 |
| 15 | 18 | 3.59 | WF2_only | WF2 | WF2 |
| 14 | 19 | 3.03 | WF2_only | WF2 | WF2 |
| 14 | 7 | 2.82 | WF2_only | WF2 | WF2 |

**Table SI 2**: Cluster-cluster interactions. Only the top 5 interactions based on probability values for each type of interaction is displayed

| | Peptide2 | WF1a | | |
|---|---|---|---|---|
| Peptide1 | Cluster1 | 4 | 5 | 7 |
| WF1a | 1 | | | 17LEU:18ASP (11.8%) -vdW<br>13GLY:18ASP (11.6%) - vdW<br>17LEU:19HIS (9.5%) vdW<br>2LEU:17LEU (8.2%) - HI<br>12ILE:18ASP (8.1%) - vdW |
| | 10 | 11ILE:14GLY (9.8%) -HI<br>10LYS:14GLY (9.5%) - vdW<br>11ILE:15ALA (8.7%) - HI<br>11ILE:13GLY (5.6%) - HI<br>11ILE:5ILE (4.9%) - HI | | |
| | 7 | | 4ARG:12ILE (18.2%) - vdW<br>1TRP:12ILE (18.2%) -HI<br>2LEU:13GLY (18.2%) - HI<br>1TRP:13GLY (9.1%) HI<br>4ARG:11ILE (9.1%) -vdW | |
| | 9 | | | 15ALA:11ILE (11.6%) - HI<br>14GLY:10LYS (11.1%) - vdW<br>14GLY:11ILE (10.2%) - HI<br>16ALA:12ILE (5.6%) - HI<br>10LYS:8GLY (5.3%)-vdW |

**Table SI 3**: Top residue-residue contacts of type WF1a:WF1a in the pure WF1a systems

| Peptide1 | Peptide2 Cluster | WF1a 10 | 12 | 7 |
|---|---|---|---|---|
| WF1a | 12 | | 18ASP:13GLY (8.7%) - vdW<br>17LEU:13GLY (6.5%)- HI<br>17LEU:12ILE (5.8%) -HI<br>19HIS:13GLY (5.8%) - vdW<br>18ASP:14GLY (5.8%)- vdW | |
| | 2 | | 15ALA:14GLY (9.6%)- HI<br>14GLY:16ALA (8.9%)- HI<br>14GLY:15ALA (8.2%) - HI<br>16ALA:13GLY (6.6%)- HI<br>16ALA:12ILE (6.0%) - HI | |
| | 7 | 11ILE:3ARG (9.9%) - vdW<br>15ALA:9VAL (9.1%) - HI<br>11ILE:1TRP (8.3%) - HI<br>14GLY:9VAL (7.2%) - HI<br>16ALA:8GLY (4.4%)- HI | | 16ALA:11ILE (14.3%) - HI<br>16ALA:8GLY (14.3%)- HI<br>15ALA:9VAL - HI (14.3%)<br>15ALA:17LEU (14.3%) - HI<br>13GLY:9VAL (14.3%)- HI |

**Table SI 4**: Top residue-residue contacts of type WF1a:WF1a in the mixed WF1a systems

| | Peptide2 | WF2 | |
|---|---|---|---|
| Peptide1 | Cluster1 | 1 | 14 |
| WF2 | 20 | 6PHE:4SER (14.3%)- vdW<br>4SER:22THR (14.3%) - HB<br>7LYS:3GLY (14.3%) - vdW<br>4SER:19ALA (14.3%) -vDW<br>6PHE:19ALA - HI (14.3%) | 24TYR:22THR (24.2%) -HB<br>25LEU:18LYS (16.2%) -vdW<br>25LEU:19ALA (15.2%) - HI<br>24TYR:24TYR (14.9%) - $\pi$-$\pi$<br>20ALA:23HIS (4.6%) - vdW |
| | 6 | 2TRP:18LYS (37.6%) -vdW<br>6PHE:19ALA (22.3%) -HI<br>7LYS:4SER (9.0%) - electrostatic<br>4SER:18LYS (6.0%) - electrostatic<br>7LYS:3GLY (4.9%) -vdW | 23HIS:5PHE (18.5%) - $\pi$-$\pi$<br>23HIS:3GLY (14.5%) -vdW<br>22THR:4SER (11.6%) - HB<br>23HIS:4SER (11.5%) - HB<br>25LEU:1GLY (8.1%) -vdW |

**Table SI 5**: Top residue-residue contacts of type WF2:WF2 in the mixed WF2 systems

| | Peptide2 | WF2 | | | |
|---|---|---|---|---|---|
| Peptide1 | Cluster1 | 18 | 19 | 4 | |
| WF2 | 14 | | 9ALA:17GLY (30.5%) - HI<br>5PHE:21LEU (29.3%) -HI<br>3GLY:23HIS (19.1%) -vdW<br>5PHE:23HIS (8.1%) -$\pi$-$\pi$<br>9ALA:20ALA (7.0%) - HI | | 18LYS:22THR (20.5%) - HB<br>19ALA:22THR (18.1%) - vdW<br>16VAL:22THR (16.9%) - vdW<br>7LYS:4SER (16.5%) - electrostatic<br>17GLY:22THR (5.7%) -vdW |
| | 15 | 21LEU:22THR (17.9%) -vdW<br>20ALA:22THR (17.3%) - vdW<br>21LEU:23HIS (16.3%) - vdW<br>20ALA:24TYR (16.1%) - vdW<br>19ALA:22THR (6.5%)-vdW | | 20ALA:22THR (17.1%) - vdW<br>21LEU:22THR (17.1%) - vdW<br>20ALA:24TYR (16.7%) - vdW<br>21LEU:23HIS (14.4%) - vdW<br>19ALA:22THR (8.5%) - vdW | |

**Table SI 6**: Top residue-residue contacts of type WF2:WF2 in the mixed WF2 systems
